# Metabolic interactions underpinning high methane fluxes across terrestrial freshwater wetlands

**DOI:** 10.1101/2024.04.15.589101

**Authors:** Emily Bechtold, Jared B. Ellenbogen, Jorge A. Villa, Djennfyer K de Melo Ferreira, Angela M. Oliverio, Joel Kostka, Virginia I. Rich, Ruth K. Varner, Sheel Bansal, Eric J. Ward, Gil Bohrer, Mikayla A. Borton, Kelly C. Wrighton, Michael J. Wilkins

## Abstract

Current estimates of wetland contributions to the global methane budget carry high uncertainty, particularly in accurately predicting emissions from high methane-emitting wetlands. Microorganisms mediate methane cycling, yet knowledge of their conservation across wetlands remains scarce. To address this, we integrated 1,118 16S rRNA amplicon datasets (116 new), 305 metagenomes (20 new) that yielded 4,745 medium and high-quality metagenome assembled genomes (MAGs; 617 new), 133 metatranscriptomes, and annual methane flux data across 9 wetlands to create the Multi-Omics for Understanding Climate Change (MUCC) v2.0.0 database. This new resource was leveraged to link microbiome compositional profiles to encoded functions and emissions, with specific focus on methane-cycling populations and the microbial carbon decomposition networks that fuel them. We identified eight methane-cycling genera that were conserved across wetlands, and deciphered wetland specific metabolic interactions across marshes, revealing low methanogen-methanotroph connectivity in high-emitting wetlands. *Methanoregula* emerged as a hub methanogen across networks and was a strong predictor of methane flux, demonstrating the potential broad relevance of methylotrophic methanogenesis in these ecosystems. Collectively, our findings illuminate trends between microbial decomposition networks and methane flux and provide an extensive publicly available database to advance future wetland research.

## INTRODUCTION

Methane (CH_4_) is a potent greenhouse gas (GHG) contributing to current atmospheric warming^1^. Despite accounting for less than 8% of the land coverage, natural wetlands represent the largest natural source of CH_4_ and contribute between 20-50% of natural global CH_4_ emissions^2–4^. Forecasting CH_4_ flux from wetlands remains challenging due to complex interactions between environmental variables such as temperature, soil moisture, and vegetation type, as well as the spatial and temporal variability of CH_4_ emissions from wetlands^3,5,6^. Furthermore, the wide array of wetland ecosystems, encompassing peatlands, marshes, swamps, and floodplains, adds complexity to the accurate quantification of CH_4_ emissions at a global scale, as each wetland potentially harbors distinct CH_4_ production processes and emission rates.

In the saturated soil conditions typical of wetlands, CH_4_ generation occurs through an interactive microbial decomposition network that hydrolyzes and ferments plant polymeric material into smaller molecular weight compounds (Figure 1B). These compounds serve as substrates for methanogenic archaea, which utilize three distinct metabolic pathways defined by their substrate preference - hydrogenotrophic, acetoclastic, and methylotrophic - for CH_4_ production^7^. Microbially derived soil CH_4_ can subsequently be emitted to the atmosphere or undergo further microbial oxidation by aerobic or anaerobic methanotrophic bacteria^8^. While this decomposition framework is well-theorized^9,10^, the extent to which these microbial members, functional guilds, and overall trophic structure are conserved across different wetlands and their relationships to CH_4_ emissions remain unclear.

**Figure 1.**
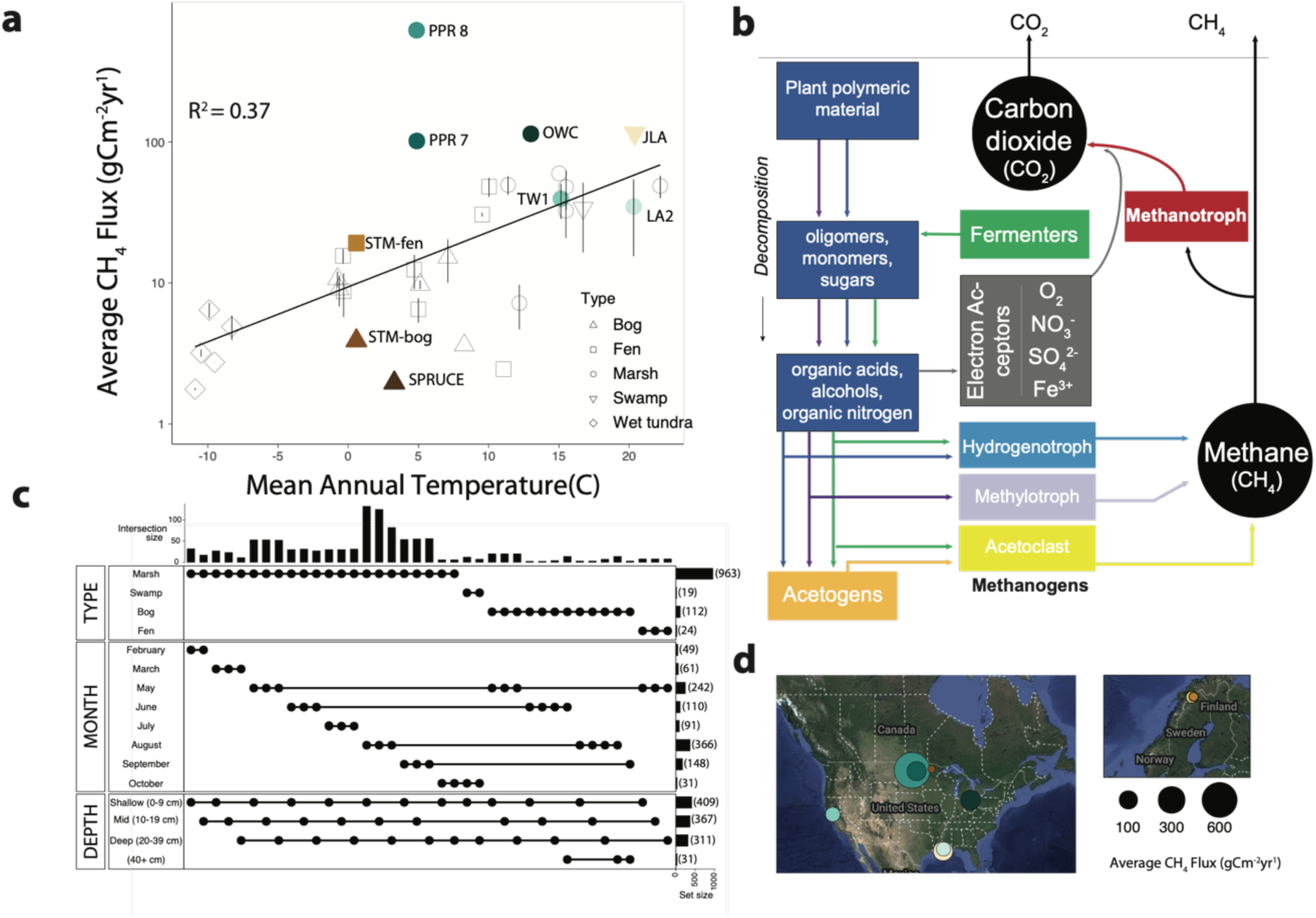
(A) Figure modified from Delwiche et al. (2021) shows mean annual methane (CH_4_) flux from wetlands included in FLUXNET-CH_4_. The deviation of the predictions from observations indicates this abiotic variable incompletely represented CH_4_ flux, especially for the highest emitting wetlands. Colored points represent sites discussed in this study. (B) Methane emissions in wetlands result from decomposition networks in which carbon decomposers first produce methanogenic substrates. These substrates are subsequently utilized by methanogenic archaea to produce CH_4_, which can either be consumed by methanotrophic bacteria or released into the atmosphere. Methanogens produce CH_4_ through three distinct pathways characterized by their substrate use: (1) hydrogenotrophic methanogenesis (reduce CO_2_ using H_2_, formate, ethanol, propanol, butanol – given in green and blue), (2) acetoclastic methanogenesis (acetate given in dark yellow or green), or (3) methylotrophic methanogenesis (CH_3_ groups cleaved from methanol or methylated amines, like trimethylamine - given in dark purple) (C) Upset plot indicates the total number of samples and their distribution across relevant categories including wetland type, sampling month, sampling depth. (D) Wetlands differ by type, annual methane flux, and geographic and climatic factors. Circle size approximates annual CH_4_. Circle area flux of proposed wetland and geographic location.

To bridge this knowledge gap, genome-resolved metagenomics has begun to unveil the identity and metabolic capabilities of microbial communities in wetland soils. This information has uncovered new methanogen and methanotroph genera^11–14^, pinpointed relevant functional pathways^15–19^, and provided insights into their spatial and temporal relevance^20^. Moreover, metagenomic data from three distinct wetlands^9,10,21^ was leveraged to construct microbial carbon decomposition networks, highlighting the microbial guilds and their constituent members involved in CH_4_ cycling withing these specific sites. While these studies laid valuable groundwork, it is imperative to complement site-specific knowledge with broader-scale analyses for a more comprehensive understanding of wetland microbiomes.

To address this broader sampling need, 16S rRNA gene amplicon sequencing characterizes bacterial and archaeal taxonomy and distribution across wetlands, albeit without providing functional content. This high throughput method allows for more extensive microbial sampling across wetland gradients, capturing microbial dynamics across wetland land coverage types, depth, and seasons^17,22–24^. Integrating knowledge from both marker gene analyses and metagenomics presents a unique opportunity to achieve comprehensive sampling of microbial conserved features, such as functional potential and network architecture across sites. Linking amplicon sequences to genomes from sampled wetland lineages would enable functional prediction, revealing the blueprints of complex wetland microbiomes at scale and transcending individual wetland boundaries.

We adopted this integrated approach for enabling genomic functional predictions for marker gene identified taxa, to uncover features of soil wetland communities and their association to CH_4_ flux across an array of freshwater wetlands. We first analyzed paired amplicon and CH_4_ flux data obtained from over a thousand samples collected across nine wetlands, representing a spectrum of CH_4_ flux rates as well as ecological and climatic conditions. From this analysis, conserved wetland-wide microbial indicators were linked to a curated genomic catalog encompassing thousands of new and existing metagenome-assembled genomes (MAGS) from wetland soils. This cross-site endeavor revealed a core set of conserved wetland microorganisms, allowing us to elucidate the functional decomposition networks supporting their activity, and delve into the physiological drivers of specific methanogenic taxa associated with high CH_4_-emitting wetlands. This study offers a comprehensive, multi-site perspective on the microorganisms and processes dictating CH_4_ dynamics in wetlands, thereby furnishing actionable insights for advancing scientific understanding and facilitating their translation and integration into climate-scale models.

## RESULTS AND DISCUSSION

### Models that rely on abiotic factors have increased uncertainty in high methane-emitting wetlands

Estimates of wetland contributions to the global methane (CH_4_) budget often rely on ecosystem-scale models, which do not represent soil microbial metabolism, but instead use abiotic variables (like mean annual air temperature) to approximate environmental states conducive for soil carbon decomposition, methanogenesis, and methanotrophy^20^. A robust meta-analysis from 42 freshwater wetlands showed that air temperature partially accounted for mean annual CH_4_ fluxes, explaining 51% of the variance across sites^25^. This discrepancy between CH_4_ flux predictions and observations for many wetlands hints at a potential role for microbial contributions in explaining these variations, a feature we sought to examine in more detail in this study.

To understand unifying microbial features across wetlands and how microbial and geochemical properties relate to CH_4_ flux, we conducted a meta-analysis using data from both published and unpublished wetland soil samples. To qualify for inclusion in our study, sites had to have amplicon sequencing data from at least 12 samples obtained from a minimum of 2 sampling depths and have CH_4_ flux measurements. From the original 42 wetlands^25^ in the noted earlier study, we identified 16S rRNA gene amplicon microbial data for three of the sites (OWC, TW1, LA2), of which the amplicon data from LA2 is newly released in this study while OWC and TWI utilize previously published data^10,27^. We also expanded the dataset to include CH_4_ flux, 16S rRNA gene amplicon, and temperature data from an additional 6 freshwater wetland sites (JLA, PPR7, PPR8, STM-fen, STM-bog, SPRUCE) (Supplemental Data 1). The incorporation of these additional sites reduced the predictive power of mean annual air temperature to explain 37% of the variability across sites (Figure 1A). Notably, the addition of sites with the highest CH_4_ fluxes (PPR8, PPR7) (Fig. 1A & 1D) reveals the limitations of mean annual air temperature as a predictor of CH_4_ flux in high emitting wetlands, such as Old Woman Creek (OWC) and those within the Prairie Pothole Regional complex (PPR).

We collated and analyzed microbial data from 1,112 samples (10% is newly released in this study) from 9 wetlands to demonstrate how incorporating knowledge of CH_4_-cycling microorganisms can contribute to improved predictive understanding of these ecosystems (Table S1, Table S2). Included data was derived from 5 marshes: Old Woman Creek (OWC), Prairie Potholes Region (PPR 7, PPR 8), AmeriFlux site US-LA2 (LA2), and AmeriFlux, site-ID US-Twt (TWI); 1 swamp: Jean Lafitte National Historical Park and Preserve (JLA); 2 bogs: Marcell Experimental Forest (SPRUCE) and Stordalen Mire (STM-bog); and 1 fen: Stordalen Mire (STM-fen). To account for inter-study variability in depth fractions, we binned these samples into three categories: shallow (0-9 cm), mid (10-19 cm), and deep (20-39 cm) (Fig. 1C).

Additionally, we supplemented these data with genomic information creating a cross-wetland genomic catalog, Multi-omics for Understanding Climate Change (MUCC) v2.0.0 database. Here we expanded the original MUCC v1.0.0 genomic catalog, which was composed of 42 metagenome and 133 metatranscriptome samples obtained from a single, high CH_4_ emitting marsh (OWC) (Figure 1A)^10^. The 2,507 medium and high-quality MAGs recovered from this wetland sampling were combined with 1,529 additional MAGS from previously published palsa, bog, and fen metagenomes from a permafrost thaw gradient at Stordalen mire (STM, Figure 1A)^9^. Additionally, we added 50 publicly available MAGs derived from the PPR complex^28^ and 43 publicly available MAGs from TWI^27^. Finally, we included 20 new metagenomes from the PPR complex, LA2, and JLA (349 Gbp of new sequencing), resulting in 617 MAGs released as new data as part of this study. In total MUCC 2.0 contains 3,634 high and medium quality, dereplicated (99% genome identity) MAGs derived from six wetland complexes totaling 8.9 Tb of sequence data (Table S3). MUCCv2.0.0 compiles previous wetland genomic datasets and expands genome representation across wetland soils spanning diverse geographies, ultimately increasing database read recruitment and reducing the computational requirements for translating reads to functional content. This wetland specific genomic resource database was used to connect microbial community profiles with functional potential.

### High CH_4_-emitting wetlands share microbial community composition and structure

Analyses across wetland sites revealed that wetland type, not geographical location, corresponded to microbial community composition and diversity. As might be expected by ecological wetland differences, bog samples derived from Sweden (STM) and Minnesota (SPRUCE), were more alike one another than bog and fen samples collected within the same wetland complex (STM). Wetlands categorized as marshes or swamps had higher bacterial and archaeal alpha diversity, higher pH, and higher CH_4_ flux than bog and fen sites (Fig. S1. Additionally, wetland type had a significant impact on community composition, and separation of communities was linked to pH (Figure 2A & S2, PERMANOVA, p<0.001). Notably, communities in bogs with the lowest pH and CH_4_ flux were most distinct from marsh/swamp communities with the highest pH and CH_4_ fluxes. Fens, with intermediate characteristics of bogs and marshes/swamps such as pH, vegetation, and nutrient levels, hosted microbial communities that were similarly intermediate of the bog and marsh communities^29^.

**Figure 2.**
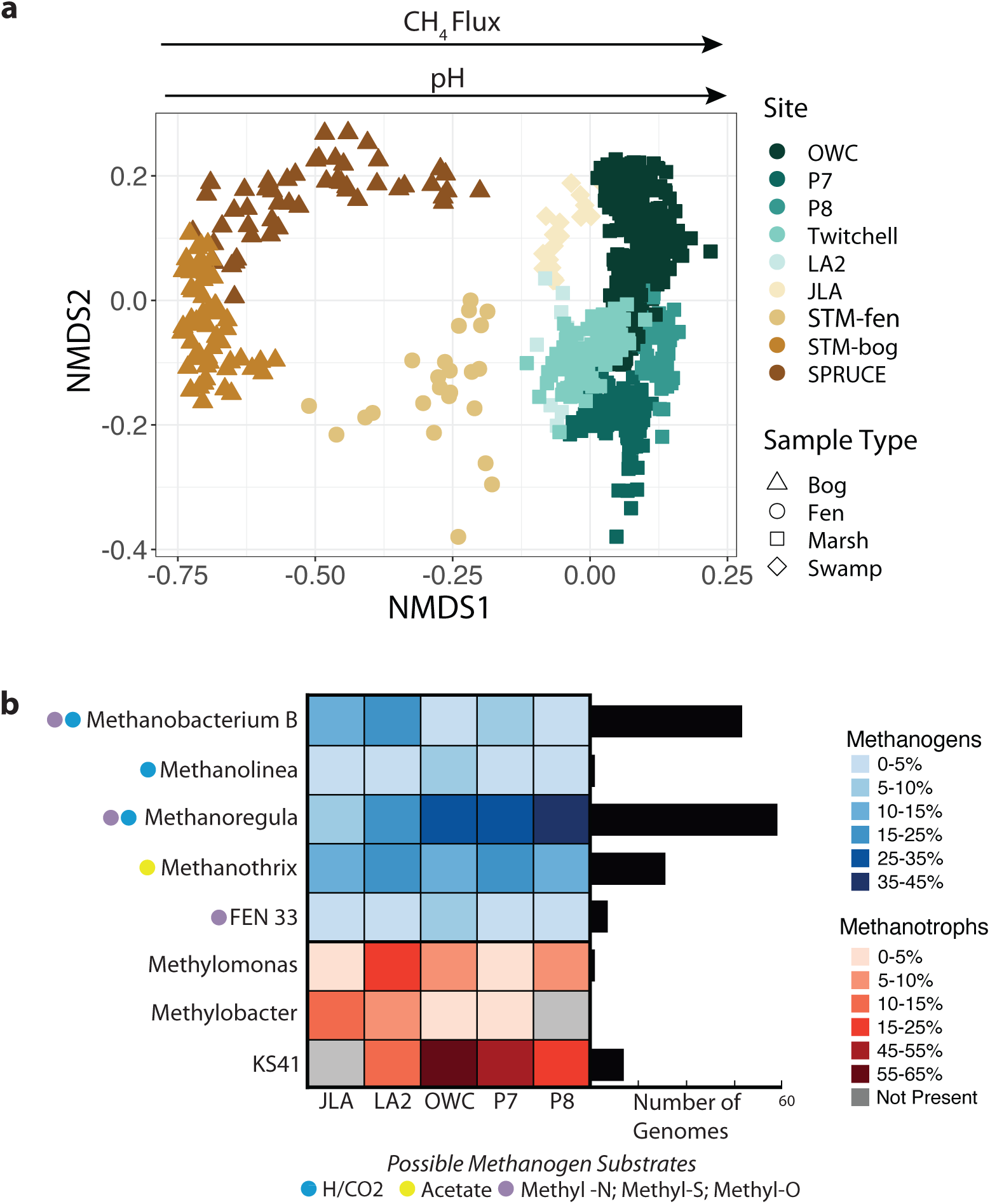
(A) Wetland type is an important control on microbiome membership and structure, despite differences in sampling strategies and geographic locations. 16S rRNA amplicon data on soil microbial communities from marsh and swamp samples cluster together (rectangles and diamonds, most right side) and are statistically distinct from fen (triangle, middle and most left side) and bog (circle, middle) microbial communities. (B) Core methane cycling members across distinct wetlands. Heatmap shows the relative abundance of each genus within the methanogen (blue) or methanotroph (red) community across wetlands. To illuminate the metabolic features of these core taxa in high methane emitting wetlands, we utilized the Multi-Omics for Understanding Climate Change (MUCC) v 2.0.0 database, with 140 MAGs assigned to our core taxa. Genome counts per genus are shown in the bar chart (black).

CH_4_ flux was loosely correlated with temperature across wetland types but this trend was absent at the level of individual wetland types. In marshes and swamps – the highest CH_4_ emitting wetland types – no correlation to temperature was observed (*R^2^*=0.17, *p*=0.16) (Fig. S3A), suggesting that other factors may be important for predicting CH_4_ flux^3,30^. We next assessed the relationships between CH_4_ flux and CH_4_-cycling microbial community members including methanogens and methanotrophs across sites. Bog and marsh sites hosted different methanogen communities (Fig. S4), with bog sites characterized by dominance of a few methanogens and low relative abundances of acetoclastic methanogens ^3,31,32^. For example, *Methanothrix*, an obligate acetoclastic methanogen was significantly more enriched in fen, marsh, and swamp samples than in bog samples. Overall, marsh and swamp sites contained a higher diversity and evenness of methanogen taxa and functional types. Collectively, the functional potential to utilize more diverse methanogenic substrates in high CH_4_ emitting marsh sites could contribute to higher CH_4_ fluxes.

To fully understand microbial contributions to the methane cycle, we also assessed the distribution of methanotroph communities across wetland types. Across all sites aerobic methanotrophs were dominant, while the anaerobic methanotrophs assigned to the genus *Methanoperedens* were enriched only in the three highest methane emitting sites (OWC, PP7, PP8) (Figure S4). We found that the diversity of methanogens (*R^2^*=0.5, *p*=0.034), but not methanotrophs (*R^2^*=0.22, *p*=0.2), was significantly correlated to CH_4_ flux (Fig S3B). Additionally, the ratio of methanogen to methanotroph relative abundances was correlated to flux (*R^2^*=0.45, *p*=0.047) (Fig S3C), but the relative abundance of methanogens and methanotrophs alone was not. This suggests the coupling of methanogens and methanotrophs act as a control over CH_4_ flux in wetland environments, highlighting how the balance between these microbial groups likely influences net methane emissions.

### Identification of a widespread, core group of CH_4_ cycling organisms

Given more consistent sampling methodology (i.e., similar sequencing protocols), as well as the higher measured CH_4_ fluxes, we focused on understanding trends in microbial dynamics across 5 marsh and swamp sites (JLA, LA2, OWC, PPR7, and PPR8) (see methods). We first assessed occupancy patterns across sites to identify if there were core methanogens and methanotrophs for these marsh samples, identifying five methanogens and four methanotrophs detected in at least one sample from each site^33^ (Fig. 2B). Despite wetland differences in site, depth, and time of year sampling (Figure 1), five core methanogen genera were found in a majority of samples: *Methanothrix* (79.7%), *Fen 33* (order *Methanomassiliicoccales*) (72.6%), *Methanobacterium B* (50.9%), *Methanolinea* (55.5%), and *Methanoregula* (93.9%). Interestingly, each methanogenic pathway (hydrogenotrophic, acetoclastic, methylotrophic methanogenesis) was represented within the core community, indicating that all three pathways are consistently important and likely utilized for wetland CH_4_ production in high emitting marsh and swamp ecosystems (Fig. 2B). Three methanotrophs were identified as core but were found in a lower percentage of samples: *Methylomonas* (60.3%)*, Methylobacter* (39.8%), and *Methylomonadaceae KS41* (85.4%). However, because the core methanotrophs require oxygen for methane oxidation, these methanotrophs may not be as detectable in the deeper anoxic samples sampled here. Constraining our analyses to only the top 10 centimeters of sediment where oxygen might be more available, we found *Methylomonas* present in 75.1%, *Methylobacter* in 57.1%, and *KS41* in 95.2% of samples. Core microbiomes have become increasingly viewed as important because of their assumed role as critical to a given ecosystems’ functioning^34,35^. Collectively, these discoveries underscore the pivotal role of select organisms in actively shaping the methane cycle within freshwater marsh ecosystems. These insights carry implications for forthcoming research activities, highlighting these organisms as candidates for more thorough physiological validation and study, as well as focus organisms for scaling to modeling endeavors.

### MUCC database enables deeper insight into trophic patterns from co-occurrence networks

For each of the 5 marsh sites, we performed network analysis based on co-occurrence patterns to help unravel possible microbial interactions within these complex, methanogen-oriented communities. We hypothesized that methanogen network structure in wetland communities would act as a predictor of CH_4_ flux. To test this hypothesis, we built 16S rRNA gene positive co-occurrence networks at each site using both the community-wide amplicon data and only the methanogen community data (Figure S5).

Although network structure of the entire community did not relate to CH_4_ flux (Figure 3K), a more constrained network comprising the significant co-occurrences that included a methanogen member did uncover important trends (Figure 3L). These networks revealed a negative correlation between the number of methanogen-related network nodes and CH_4_ flux, indicating a relationship between less complex methanogen networks and higher annual CH_4_ emissions. Furthermore, the number of methanotrophs associated with methanogens in these networks was greater in the lower methane emitting sites (JLA, LA2), indicating that lower CH_4_ fluxes are associated with communities where methanotrophs and methanogens co-occur. In contrast, while high CH_4_-emitting sites (OWC, PPR7, PPR8) host methanotrophs and methanogens, they were generally linked by fewer connections (Figure 3M). Methanotrophs can act as a filter, oxidizing anywhere from 20-60% of the CH_4_ before it is released into the atmosphere^3,36,37^ and these results indicate that their absence in wetland samples where methanogens are present could contribute to greater CH_4_ fluxes.

**Figure 3.**
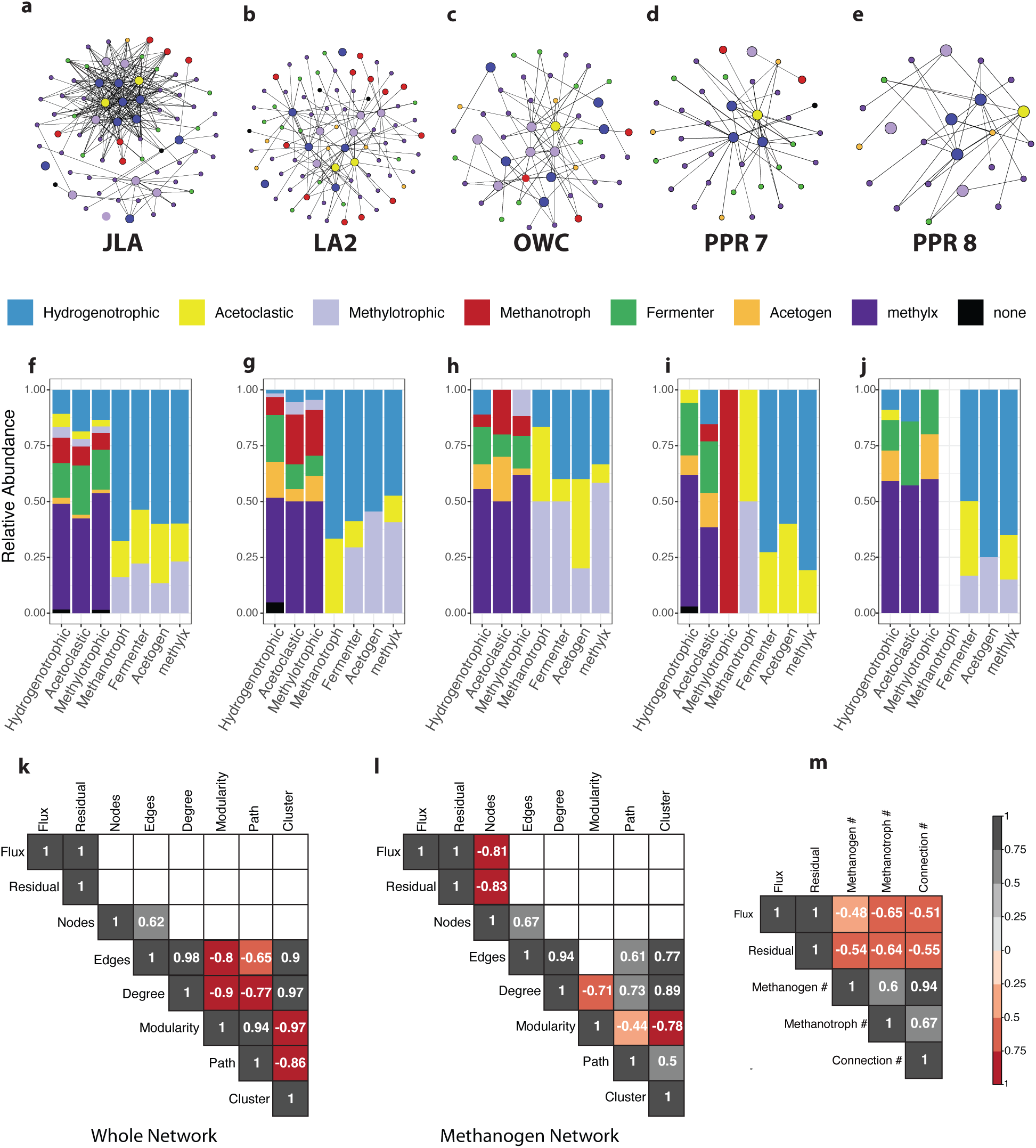
(A-E) Co-occurrence network analysis revealed network structure of methanogen associated taxa across wetlands. Networks depicting site specific co-occurrence analysis uncovered the network of microorganisms coordinated to methanogens across each site, with nodes representing microbial taxa. Larger nodes represent methanogens, while small nodes represent bacterial taxa. Nodes are colored by inferred metabolic potential of 16S rRNA linked MAGs within MUCC. (F-J) Proportion of connections between groups in each network are given in the bar charts below and show conserved patterns in network connections across sites. Missing bars indicate no connections. Correlation between network statistics and methane flux measurements derived from the Ameriflux network was measured for (K) whole community networks and (L) methanogen networks. Only number of nodes in the methanogen network was correlated with methane flux. (M) Additionally, negative correlation between annual methane residual and methane flux (from Figure 1) to number of methanogens, methanotrophs, and connections between the two were observed.

To determine potential metabolic interactions that underpin CH_4_ production across these sites, we developed metabolic profiles for methanogen-connected taxa in our 16S rRNA gene networks. Utilizing the MUCC 2.0.0 database, we linked microbes present in the networks with MAG representatives and assigned them functional categories: obligate fermenter, homoacetogen, demethylating, or none of these three criteria (Fig. 3A-E, 4 & Table S4. We selected these criteria, as they are thought to cross feed methanogens (Figure S1, Data Table) and are traits that can be inferred from genomes clearly. Methanogen networks were composed of 699 unique co-associated genera, of which 131 genera had a genome representative in the MUCC database (Figure 4). Summarizing these genome representatives within the methanogen networks, 12 were categorized as methanogens, 7 as methanotrophs, 23 as obligate fermenters, 8 as homoacetogens, 1 as both obligate fermenter and homoacetogen, and 75 demethylating (methyl-x), and 4 did not meet these criteria (Rules for assignment are found in Table S4). Additionally, 6 methanogens and 10 methanotrophs identified based on 16S rRNA gene taxonomy alone (no matches to MUCC, but metabolism is defined in literature) were included in the networks (Fig. 4, Table S5).

**Figure 4.**
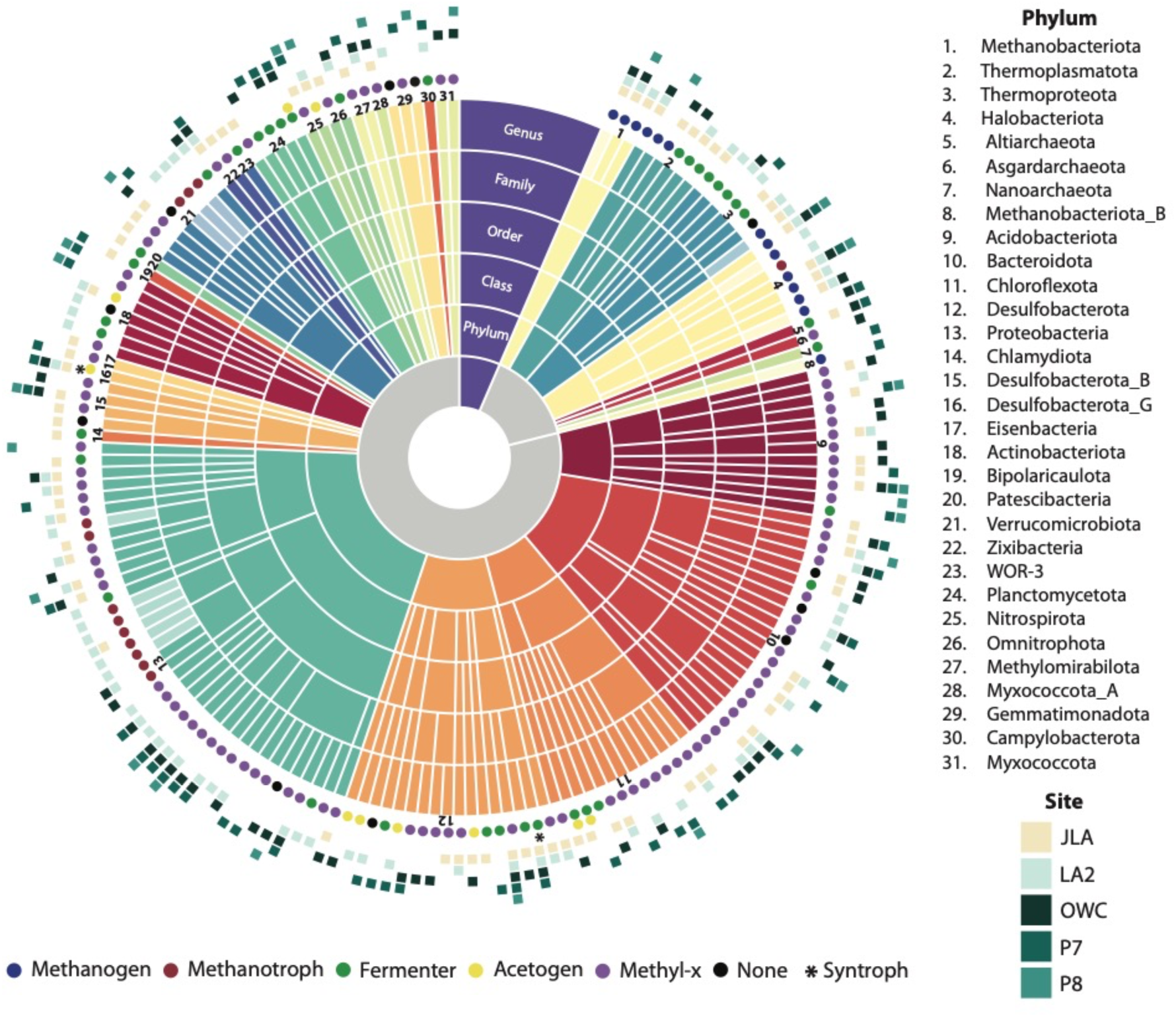
Taxonomy of the 158 genera represented in the networks that are found within the MUCC database. Additionally, 6 methanogens and 9 methanotrophs were identified based on 16S rRNA were included in the networks and are shown in the network with reduced opacity at the genus level. Circles around the edge represent inferred metabolic potential and squares represent the sites where the genus had significant co-occurrence with a methanogen.

Specifically, obligate fermenters have the potential to produce acetate, formate, and H_2,_ which we hypothesized would directly promote methanogen activity^38,39^ and thus be positively associated with our methanogen networks. As we expected, obligate fermenters were highly connected to hydrogenotrophic and acetoclastic methanogens, likely supporting cross feeding. In total, obligate fermenters had 99 significant interactions with methanogens of which 73% were to hydrogenotrophic or acetoclastic methanogens (Fig. 3F-J). Additionally, obligate fermenters were found to highly co-occur with certain methylotrophic methanogens such as *Methanofastidiosum*, which requires H_2_ to reduce methylated thiol to form methane. Compared to hydrogenotrophic methanogenesis, this form of methanogenesis is more thermodynamically favorable under low H_2_ conditions and has been proposed to support H_2_ producing syntrophs and fermenters by preventing accumulation of H_2_^12^. In summary, anoxic carbon exchanges between obligate fermenters and methanogens appear vital to carbon cycling in wetlands.

Syntrophy denotes a symbiotic interaction among diverse microorganisms, wherein the exchange of metabolic byproducts mutually supports each organism’s metabolism. This phenomenon is particularly prominent in methanogenic environments, where methanogens play a crucial role in regulating product concentrations, thereby rendering otherwise endergonic processes thermodynamically favorable^40^. In our study, we investigated obligate fermenters to uncover evidence of secondary fermentative syntrophs, identifying two prevalent syntrophic genera across methanogen networks: *Smithella*, present in four marshes except PPR8, and *Syntrophorhabdus*, found across all five marsh networks. Previous research has demonstrated the capacity for acetate and hydrogen production by Syntrophorhabdus, aligning with our genome-based characterization of these 7 MAGs in MUCC. Notably, in our networks, Syntrophorhabdus exhibited multiple (8) connections to hydrogenotrophs and acetoclasts, further emphasizing its role in metabolic exchanges. These genomic metabolic insights highlight the intricate connections harbored within these co-association networks, exchanges essential for maintaining metabolic efficiency in methanogenic environments.

Homoacetogens are also interacting with methanogens, as these microorganisms grow on H_2_/CO_2_/CO and produce acetate as the main metabolic product. We hypothesized that these organisms could cross-feed acetoclastic methanogens^15^ and or could compete with hydrogenotrophic methanogens for substrates^41^. The 9 homoacetogen MAGs identified in the methanogen networks comprised 15 nodes and were closely related across sites, belonging to two main phyla, Desulfobacterota and Chloroflexota despite many other acetogens across other phyla existing in the MUCC database. We observed 32 associations between these acetogens and methanogens, with 50% to hydrogenotrophic, 28% to acetoclastic and 22% to methylotrophic methanogens. Additionally, 6 of the 8 acetoclastic methanogens had at least one connection to an acetogen, supporting our hypothesis that acetogens were cross-feeding methanogens. While our finding does not preclude competition between hydrogenotrophs and other acetogens, these identified positive associations may reflect sufficient hydrogen production within the soil profile to support co-existence of both guilds, or the separation of guilds across microsites.

Finally, demethylating microorganisms, whether bacteria or archaea, are capable of removing methyl groups from oxygen, sulfur, and nitrogen (O, S, N) containing compounds. Unlike methylotrophic methanogens, these taxa do not produce methane directly; however, they may engage in cross-feeding or competition dynamics with methylotrophic methanogens. Depending on the enzymatic systems they encode, these microorganisms can lead to several outcomes: (i) production of trimethylamine (TMA), a substrate for certain methanogens; (ii) formation of quaternary amines (QA), which can could be utilized by select methylotrophic methanogens; or (iii) direct utilization of methylated O, N, or S compounds, which may (iiia) compete with methylotrophic methanogens or (iiib) generate acetate and hydrogen to support hydrogenotrophic or acetoclastic methanogens. The methyl-metabolism category exhibited substantial connectivity with methanogens, comprising nearly half of the connections across sites. Notably, 68% of these connections (comprised mostly of type iii demethylating microorganisms) were linked to acetoclastic and hydrogenotrophic methanogens not methylotrophs suggesting that demethylating metabolisms in soils could indirectly bolster non-methylotrophic methane production. These findings underscore the complexity of microbial interactions beyond methane production and oxidation, thereby contributing to a more comprehensive understanding of microbial cross-feeding and its broader implications for methane emissions.

### *Methanoregula* is critical for CH_4_ production in wetlands

Two core methanogens (Figure 2), *Methanothrix* and *Methanoregula*, were found in networks across every marsh indicating global importance in the wetland CH_4_ cycle. *Methanothrix* is an obligate acetoclastic methanogen already shown to be globally distributed and an important contributor to CH_4_ emissions in wetlands^16^. *Methanoregula* has been found in wetlands and other habitats around the world, and like at many of our sites, is a prominent member of methanogenic networks and consistently a dominant methanogen^42,43^. We found that its dominance (proportion of methanogens that are *Methanoregula*) was related to CH_4_ flux, such that percent of methanogens that are *Methanoregula* significantly correlated to CH_4_ flux and the residual values that were not well predicted from the temperature-CH_4_ flux correlation in Figure 1 (Figure 5A). Additionally, we tested how well temperature, *Methanoregula* dominance, and the two combined explained methane flux. When looking at the 9 study sites, CH_4_ flux was not predicted by temperature alone (*R^2^*=0.15, *p*=0.30,), was predicted by *Methanoregula* dominance (*R^2^*=0.54, *p*=0.02,), but that temperature combined with *Methanoregula* dominance was the best predictor (*R^2^*=0.84, *p*=0.02). This is one example of how incorporating biological insights with already existing abiotic data could improve the predictive power of climate models.

**Figure 5.**
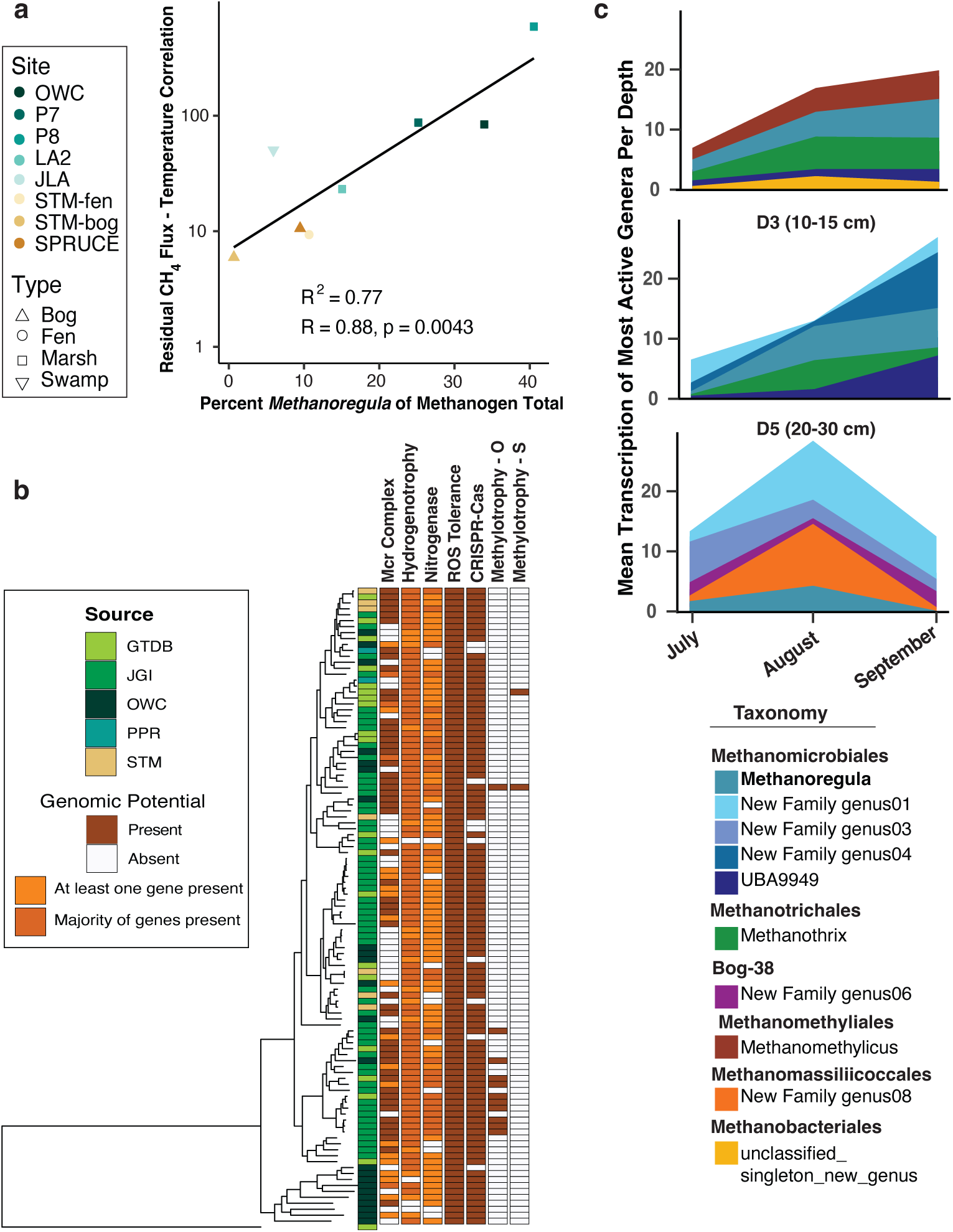
(A) Residual values from the methane flux to temperature trend line was significantly related to the relative abundance of *Methanoregula* within the methanogen community. (B) Genome tree of Methanoregula MAGs from MUCC (OWC, PPR, STM), plus available MAGS from JGI and GTDB. A pangenome-analysis shows the largely conserved encoding of genes for key physiological features, as well as limited novel metabolic potential (e.g., methylotrophic genes) which may directly or indirectly support high methane fluxes from *Methanoregula* in wetlands. (C) Mean transcription of top five most active methanogenic genera at three depths (0-5 cm, 10-15 cm, 20-30 cm) in the mud site type across the 2018 sampling season. predictive of CH_4_ fluxes.

To understand potential physiological drivers that link *Methanoregula* and predications of CH_4_ flux, we conducted a genomic analysis of 107 dereplicated MUCC-derived and publicly available (i.e., GTDB, JGI) MAGs. *Methanoregula* encoded diverse metabolic strategies, the capacity for fixing nitrogen (nitrogenase), viral defense (CRISPR-Cas), and mechanisms to respond to fluctuating redox conditions (reactive oxygen species) (Figure 5B). *Methanoregula* are classically designated hydrogenotrophic^44^, which we broadly confirmed here (Figure 5B). We also report that some *Methanoregula* genomes encode genes for methylotrophic methanogenesis, specifically for the demethylation of methylated sulfides^45^ and methoxylated^19^ compounds, compounds prevalent in wetlands^10,15^. Although hydrogenotrophic methanogenesis is generally recognized as the dominant CH_4_ -generating pathway in wetlands, recent studies have indicated that methylotrophic methanogenesis contributes more to CH_4_ flux than previously realized^17,21,46^. Therefore, the apparent significance of *Methanoregula* in contributing to CH_4_ emissions across diverse wetlands and within wetland gradients could partly be explained by a broader than previously understood ecological niche.

To investigate the role of *Methanoregula* within a high CH_4_ emitting wetland, we mined a previously undefined role for *Methanoregula* from 39 paired metatranscriptome and metabolome datasets across spatial and temporal gradients from a single mudflat at OWC^10^ (Figure S6A). At this mud-type site, a *Methanoregula* MAG (OWC-0053) was one of the transcriptionally most active methanogens throughout the entire soil column across 3 months of peak CH_4_ production (Figure 5C). This genome was also one of the 9 genomes that predicted 78% of soil porewater CH_4_ concentration (Figure S6B). In summary, our comprehensive analysis reveals *Methanoregula’s* substantial contribution to CH_4_ dynamics within a high-emission wetland, highlighting its prominent role as a key player in CH_4_ production across spatial and temporal scales.

These findings help in part explain the significant correlation between *Methanoregula* abundance and CH_4_ flux across wetlands, and its role in marsh CH_4_ networks. Our results suggest that *Methanoregula* may possess a broader physiological capacity to produce CH_4_ and adapt to various abiotic and biotic constraints present in marsh soils. By shedding light on the functional significance of *Methanoregula*, a core taxon across wetlands, our study contributes to advancing our understanding of wetland CH_4_ emissions. Our findings use a cross-site analysis to identify core lineages, like *Methanoregula*, warranting further physiological exploration, as the metabolic assumptions may be constrained by prior strict substrate and redox capabilities. Ultimately our results show promise for biological knowledge to enhance predictive models of wetland emissions, ultimately facilitating more effective management and mitigation strategies.

## Conclusions

Microbial processes related to CH_4_ flux have been well-characterized at a handful of individual sites. However, site-specific knowledge of wetland microbiomes suffers from limited generalizability, as wetland ecosystems vary widely. Therefore, insights gained from studying microbiomes in one wetland may not necessarily apply to others, restricting the broader understanding of wetland microbial communities and their roles in ecosystem processes. Here, we build on existing single-site studies by building a multisite wetlands database, and synthesizing decomposer and CH_4_-cycling networks and their relation to CH_4_ flux data across multiple wetland ecosystems. Linking 16S rRNA gene data to genomes from the MUCC database, we developed metabolic profiles for methanogen-connected taxa. We found microbial cross-feeding has broad implications for CH_4_ emissions across wetland environments. Additionally, the highest CH_4_ emitting wetlands had the fewest methanogen network connections, suggesting streamlined metabolic circuits may contribute to enhanced CH_4_ production across wetland soils. Finally, we revealed that *Methanoregula* is a key contributor to CH_4_ flux in wetland environments, potentially due in part to previously unknown metabolic versatility. Ultimately, MUCC is a powerful microbiome tool enabling us to decode microbial organismal and metabolic patterns across multiple environments, with the goal of improving predictive modeling frameworks.

## Methods

### Multi-Omics for Understanding Climate Change (MUCC) v2.0.0 Database

Data was compiled from 9 different wetlands (5 marshes, 1 swamp, 1 fen, and 2 bogs), including both previously published and unpublished datasets. Published data were sourced from Old Woman Creek (OWC), AmeriFlux site-ID US-Twt (TWI), and SPRUCE; both published and unpublished data was compiled from Prairie Potholes Region (PPR 7, PPR 8) and Stordalen Mire (STM-fen and STM-bog); and unpublished data were collected from Jean Lafitte National Historical Park and Preserve (JLA) and AmeriFlux site US-LA2 (LA2). The Multi-Omics for Understanding Climate Change (MUCC) v2.0.0 database combines 997 16S rRNA, 284 metagenomic, and 133 metatranscriptomic datasets from PPR, STM, OWC, TWI, and SPRUCE, along with 115 newly analyzed 16S rRNA and 20 metagenomic samples from PPR, JLA, and LA2. DNA extraction and amplicon sequencing info for all sites can be found in Table S7. Accession numbers for all samples can be found in Table S1, while sample IDs and GTDBk v207 taxonomy for 16S rRNA data are in Table S2, and the details of 4,745 medium and high-quality Metagenome Assembled Genomes (MAGs) are listed in Table S3. The MAGs and 16S rRNA data from MUCC v2.0.0 are available on Zenodo (https://zenodo.org/records/10822869) and NCBI (PRJNA1007388).

#### Old Woman Creek (OWC)

OWC National Estuarine Research Reserve (41° 22’N 82°30’W) is located on the southern shore of Lake Erie in Ohio. It is composed of a permanently flooded channel surrounded by marsh, occasional mud flats (which are inundated most of the time), and an upland forested habitat^16^. In brief, sediment cores were collected from sites representing distinct eco-hydrological patch types (cattail plant, mud, and open water) in triplicate in May, June, July, August, and September of 2018 using a modified Mooring System soil corer^16^. Cores, sampled to a depth of 35cm, were sub-sectioned into six depths using a hydraulic extruder: 0-5 cm, 5-10 cm, 10-15 cm, 15-20 cm, 20-25 cm, 25-30 cm. Microbiome data from 626 samples included bacterial and archaeal 16S rRNA amplicon sequence data, metagenomes, and metatranscriptomes^10,16^. Meteorological and eddy-covariance flux data for the site are available through AmeriFlux, site-ID US-OWC^47^. Gap-filled and averaged data used in this analysis were obtained from FLUXNET-CH4^30^.

#### Prairie Pothole Region (PPR)

Cottonwood Lake Study Area (47° 05’N: 99° 06’W), located northwest of Jamestown, North Dakota, is a protected area owned by U. S. Fish and Wildlife Service and is a long-term research site (>30 years) for the U. S. Geological Survey (USGS). The 92-ha site consists of 17 distinct wetlands with permanent-to-temporary inundation. Samples were collected from two permanent wetlands: P8 (47° 05’55.8”N 99°06’14.1”W) and 2 sub-locations within P7 - Location 1 (47°05’43.7”N 99°06’00.8”W) and Location 2 (47°05’46.7”N 99°05’57.9”W). Cores were collected in triplicate at each location in March, May, and September of 2015 using a modified Mooring System soil corer. Cores, sampled to a depth of 30 cm, were sub-sectioned using hydraulic extrudation in 3-cm increments. MUCC v 2.0.0 included 214 16S rRNA sequencing samples and 18 previously published metagenomes^24^ combined with 18 new metagenomes from PPR.

Annual CH_4_ flux data was averaged from 2011-2016^48^. Methane fluxes were measured using the static chamber method^49^ every two weeks during the growing season (defined as soil temperature ≥5 °C). During each sampling event, chambers were floated in open waters of P7 and P8 for 30 minutes after which headspace gas samples were collected through a rubber septa and stored in evacuated 10-ml serum vials. Sample gases were analyzed for methane concentrations on a gas chromatograph equipped with electron capture and flame ionization detectors (SRI Model 8610, California) located at the USGS Northern Prairie Wildlife Research Center. Methane flux rates were calculated using the linear change in CH_4_ concentration during the deployment, chamber dimensions and temperature, and the Ideal Gas Law. Biweekly flux rates were scaled to annual cumulative CH_4_ flux by summing the mean flux rates between consecutive sampling events and multiplying by the time between events.

#### Louisiana Wetlands (JLA and LA2)

Two distinct sites were sampled in Louisiana in October 2021. Jean Lafitte National Historical Park and Preserve (JLA) (29°80’18″ N 90°11’02″ W) and AmeriFlux site-ID US-LA2^50^ (LA2) (29°51’31.4″ N, 90°17’11.3″ W) on the Salvador Wildlife Management Area are located in coastal Louisiana. The JLA wetland is a Cypress-Tupello swamp with distinct hollow and hummock features, and the LA2 wetland is a fresh flotant marsh vegetated by a mix of *Typha sp.* and *Sagittaria sp*. In JLA, triplicate soil cores were collected using a Russian Peat Corer, and 0-10 cm and 30-40 cm intervals were sampled. In LA2, triplicate slurry samples from 0-10 cm and 20-30 cm were collected using a sipper.

Samples were kept on dry ice after processing. DNA was extracted using Zymo Research Quick-DNA™ Fecal/Soil Microbe Microprep Kit, following the manufacturer’s protocol. Amplicon libraries were prepared using a single step PCR to amplify the V4 region of the 16S rRNA gene with the primers 515F/806R ^51^ following the Earth Microbiome Project (EMP) PCR protocol. Pooled DNA products were sequenced on the Illumina MiSeq Platform using 251 bp paired-end sequencing chemistry at the Microbial Community Sequencing Lab (University of Colorado Boulder).

Gap-filled and averaged flux data for LA2 that were used here, were downloaded from FLUXNET-CH4^30^, while JLA flux was measured in four field campaigns in June, August, October, and December of 2021. Measurements were conducted using a trace gas analyzer (LICOR 7810) coupled to a custom-made chamber in triplicate 2-minute deployments in three hollow and three hummock locations. Flux was calculated following procedures described in Villa et al. 2021^52^.

#### Twitchell

Twitchell Island (121.65°W, 38.11°N), located in the Sacramento-San Joaquin River Delta, CA, hosts a USGS wetland restoration site. Meteorological and flux data for the site are available through AmeriFlux, site-ID US-Twt^53^. The Twitchell experimental wetlands are categorized as freshwater marsh. All data used from the Twitchell site were previously published in He et al^27^. Flux data was downloaded from FLUXNET-CH4^30^.

#### SPRUCE

The SPRUCE experiment (47°30.4760N; 93°27.1620W), located in the S1 bog of the US Department Agriculture (USDA) Forest Service’s Marcell Experimental Forest, is located northeast of Grand Rapids, Minnesota. All data used was published in Wilson et al 2021^22^. In this study, only data from samples collected from +0 and ambient treatments were used.

#### Stordalen Mire (Stm)

Stordalen Mire (0°34’25.7”N; 37°34’30.1”E) located near Abisko, Sweden is an Arctic permafrost peatland that covers three main habitats across a discontinuous thaw gradient: palsa, bog, and fen. Palsa overlays intact permafrost and is well-drained and dominated by woody and ericaceous shrubs. Bog overlays partially thawed permafrost, with a perched water table and *Sphagnum* moss dominance. Fen is fully thawed, inundated, and sedge-dominated. The Mire was surveyed in 2015 at a range of distributed palsas, bogs and fens; only bog and fen 16S rRNA gene amplicon data are used in this study. A serrated knife was used to cut vertically into the peat, and microbial samples were collected to fill 2ml Eppendorf tubes from each depth: “shallow” (median of 2cm, range 1-3cm); “middle” (median of 12cm, range 10-12cm); and “deep” m(edian of 20cm, range 18-20cm). Sample tubes were stored on ice in the field and transferred to -80C within 10 hours of collection. DNA was extracted with the PowerSoil 96-Well Soil DNA Isolation kit (MO BIO cat# 12955-4) following the manufacturer’s protocol. 16S rRNA gene amplicon sequencing were performed by Argonne National Laboratory using the Earth Microbiome Project barcoded 515F-806R primer set and protocol, and on an Illumina MiSeq sequencer. MAGs from 214 previously published metagenomes were also used^9^. Methane flux data for Stordalen bogs and fens were annual averages from 2012-2018 of autochamber measurements (static, closed systems) that include three replicate measurements per cover type^54^.

### 16S rRNA Gene Sequencing and Analysis

All raw amplicon sequence data were processed using the QIIME2 (v2021.2) pipeline^55^. Data from OWC, PPR, LA2, JLA, STM-f, STM-b, and Spruce sites were independently processed through QIIME2 to account for sequencing run biases. Datasets were uniformly trimmed to the same length (195 bp), paired end read were merged, and ASVs assigned using the naïve Bayes sklearn classifier trained with the GTDB-Tk (v2.1.1 r207)^56^, prior to merging at the ASV level across datasets. Because Twitchell was sequenced using a different primer set, sites were merged at the genus level. Due to a wide range in sequencing depth across sites, all samples were rarefied to 5000 reads resulting in a final dataset of 1118samples (Figure 1C). 43 samples were not retained because they fell below the minimum read depth. Across the 9 wetlands included in this study, core depth and interval sections varied. The compiled studies had different depth thresholds used to categorize shallow, middle, and deep sediments. To standardize depth measurements, we created 3 categories that encompassed the categories across studies: shallow included samples in the 0-9 cm horizon, middle included samples collected from 10-19 cm, and deep for samples collected from 20-40 cm.

### Genome assembly and binning

Previously published metagenomic samples were combined with newly analyzed samples in this release of MUCC. 20 newly analyzed samples contributed 617 MAGs (Table S1 & S3). MAGs were recovered from:

1. 2021 LA Field Sample (n=1)
2. 2021 JLA Field Sample (n=1)
3. 2022 PPR Field Sediment Samples (n=7)
4. 2022 PPR Field Water Samples (n=2)
5. 2022 PPR Lab Enrichment Samples (n=9)

LA and JLA metagenomes were processed separately from the PPR metagenomes. Raw metagenomic reads were trimmed using Sickle (pe)^57^ and assemblies were generated using Megahit (v1.2.9)^58^ with parameters --k-min 31 --k max 121 --k-step 10. Subsampled assemblies using 25% of sequencing reads were generated using IDBA-UD v 1.1.3^59^ with default parameters. Reads were mapped to contigs greater than 2500 bp using BBMap (v 38.89)^60^ and were subsequently binned using MetaBAT2^61^. Only medium and high-quality bins based on adapted MIMARKS standards (completeness >=50% and contamination <10%) were retained^62^. PPR bins from these assemblies were combined with bins from metagenomic assemblies derived from earlier sampling of PPR^28^, were combined with the bins from LA2 and JLA, and with publicly available bins from OWC^10^, STM^9^, and TWI^27^. This bin pool was dereplicated using dRep (v 3.0.0)^63^ at 99% identity. MAG completeness and contamination was estimated using CheckM^64^ and taxonomy assigned using GTDB-tk v2.3.0 with GDTB database release 207^56^.

### Community Analysis

To determine the extent to which microbial community structure varied with both wetland type (marsh, swamp, fen, bog) and sample depth (shallow, mid, deep), we conducted permutational analysis of variance (PERMANOVA) using Bray-Curtis distances. Results were visualized using non-metric multidimensional scaling (NMDS). PERMANOVA and NMDS were conducted using the vegan package ^65^ and visualized using *ggplot2*^66^ in R Studio^66^. We also correlated environmental parameters including pH, mean annual temperature, mean annual precipitation, latitude, longitude, and CH_4_ flux with microbial community structure using the R-function *envfit* (as visualized in Figure S2). Alpha diversity of the entire microbial community, of methanotrophs and methanogens, of the methanogens only, and of the methanotrophs only was calculated using the Shannon diversity index. Differences in alpha diversity between bogs and fens were calculated using analysis of variance (ANOVA). Marshes and swamps were grouped together because they have similar characteristics to each other such as pH while bog and fen were grouped because they are both types of peatland characterized by low pH and occur in similar climates^67^. Shannon diversity was correlated with individual environmental parameters using a linear regression and *corrplot* in R. Linear models were used to assess if mean annual temperature (MAT) and/or relative abundance of *Methanoregula* was predictive of methane flux across wetlands using the *lm* function in R. MAT and *Methanoregula* relative abundance were also individually tested using a regression model conducted using the R-function *ggpubr*^68^.

To determine if certain methanogens and methanotrophs were widespread (found across all sites) or restricted to specific wetland types (i.e., marsh), we conducted a core community analysis. This analysis was conducted across all samples both regardless of sample depth, and within the depth categorization to understand if core members are more likely to be present in different depth zones. Because of the wide range of sampling schemes across sites, a microbe was determined to be a core member if it was present across all sites or all sites within a categorization (marsh/swamp or bog/fen). Core analysis was preformed using ‘*summarise*’ and ‘*filter*’ commands in Tidyverse^69^.

### Co-occurrence Networks

To understand if co-occurrence patterns related to methane flux, we created co-occurrence networks based on the entire community and significant co-occurrence patterns with methanogens from JLA, LA2, OWC, PPR 7, PPR 8. We focused on these five marsh sites because we were interested in patterns within the highest methane producing communities and because these all used the same amplicon primers. Because networks are sensitive to number of input samples, each individual site’s network was composed of 12 different community samples that were randomly sampled. Additionally, samples all came from similar points in the season (September or October) and represented all sampling depths.

Network analyses were carried out in R using the packages igraph^70^, Hmisc^71^, and Matrix^72^. To determine co-occurrence patterns in the microbial communities, we used rarefied genus tables. Genera with less than 10 read counts were removed from the analysis. We used Spearman correlations to determine if genera were significantly correlated with a p-value cutoff of < 0.05 and rho of > 0.5. Gephi (0.10.1)^73^ was used to visualize networks and calculate network parameters including number of edges, nodes, average degrees, average path length, and modularity. Network parameters were correlated to methane flux using *corrplot* and linear regressions in R. Given our interest in the metabolic interactions of microbial taxa with methanogens, we focused downstream analyses on positive interactions.

To uncover the metabolic interactions patterns of the methanogens, co-occurrence networks were compared to MAGs in the MUCC database that had been assigned taxonomy using GTDB-Tk (v2.1.1 r207)^56^. Every MAG that appeared in the methanogen networks (determined if MAG and 16S ID matched at the genus level) were compiled and annotated using DRAM (v1.4.4)^74^. MAGs were further physiologically curated using DRAM curations and manual analyses, and subsequently put into one of the following categories: Methanogen, methanotroph, fermenter, acetogen, methyl-x, or other (Table S5). Methanogens, methanotrophs, and fermenters were defined using the rules set published in Olivero et al^10^. Additional methanogens and methanotrophs were assigned if a MAG was not present for that genus but has been recognized in the literature. Acetogens were assigned if they had at least 6 out of 10 steps of the Wood-Ljungdahl pathway. Methyl-x were assigned based on the presence of known substrate-specific methylotrophic genes including both aerobic and anaerobic metabolisms. All rules are outlined in Table S4. If multiple MAGs existed for each genus, over 50% of the MAGs had to follow the rules laid out above for it to be classified within a given category.

### Phylogenomic and physiological analysis of *Methanoregula*

MAGs in the MUCC database were taxonomically assigned using GTDB-Tk (v2.1.1 r207)^56^ and *Methanoregula* MAGs (n=37) were parsed by genus from the full database. Further, publicly-available *Methanoregula* MAGs were retrieved from GTDB (n=21) and JGI (n=91). These 149 MAGs were dereplicated at 99% using dRep^63^ in 107 representative MAGs. All MAGs were annotated using DRAM (v1.3.2) ^74^.

Phylogenomic analysis of the 107 dereplicated *Methanoregula* MAGs was performed using GTDB-Tk v2.1.1 r207^56^ run using the de novo workflow. The alignment was based on 53 concatenated archaeal marker genes, and a GTDB-derived genome from the phylum Undinarchaeota (GCA_002495465.1) was used as an outgroup to root the tree. The generated tree was read and visually modified, including the representation of physiological potential, in R using the ggtree package^75^. Newick tree is available at https://zenodo.org/records/10822869.

*Methanoregula* MAGs were screened for physiological potential for methanogenesis (*mcrABG*), hydrogenotrophy (genes encoding the Wood-Ljungdahl pathway), nitrogen fixation (nitrogenase) and CRISPR-Cas associated proteins using DRAM. Meanwhile, to search for possession of genes encoding reactive oxygen species (ROS) detoxification enzymes, MAGs were searched via BLAST-P using a FASTA reference file (https://zenodo.org/records/10822869) of Uniprot and KEGG-derived reference sequences of ROS detox enzymes methanogens are known to encode^76^.The BLAST-P output was limited to include only hits with both a bitscore of ⦥100 and ⦥30% identity to the target sequence. Last, to curate methylotrophic potential, we carried out the strategy used by Ellenbogen *et al.*^15^. MAGs were searched via BLAST-P using a FASTA reference file^15^ of known methylotrophic genes, namely those encoding substrate-specific corrinoid-dependent three component methyltransferase systems comprised of a substrate:corrinoid methyltransferase, a corrinoid-binding protein, a methylcorrinoid:carbon-carrier methyltransferase, and a reductive activase. The BLASTP output was limited to only include hits with a bitscore >60, and only genes from MAGs found to possess genes for directly substrate-interacting substrate:corrinoid methyltransferases were retained. Genes meeting these criteria were phylogenetically analyzed using ProtPipeliner to build RaxML trees (https://github.com/TheWrightonLab/Protpipeliner/blob/master/protpipeliner.py) relative to reference genes including those used in the BLAST-P search, plus other homologous sequences derived from UniProt from physiologically characterized methylotrophic methanogens and acetogens (Table S2 tab FASTA_reference_for_genes_trees). Newick trees are available at https://zenodo.org/records/10822869. Trees were visually inspected in iTOL^77^, and tree placement – plus gene synteny, as methylotrophic genes are often co-encoded^78,79^ - was used to confirm or refine the specific identification of genes.

### Metatranscriptomic analyses

Metatransciptome analyses was performed using a previously published normalized read count table^10^. In brief, raw metatranscriptomic reads were quality trimmed, mapped to MUCC v 1.0.0, per gene read counts were estimated, and resulting read counts were normalized to gene length and TMM normalized using log2 normalization^80^. Mean geTMM values for all genes were summed for each MAG, to generate a total expression metric for each MAGs activity within the 2018 OWC metatranscriptomes. Only metatranscriptome data from mud type sites are included in these analyses. These MAG totals were further summed to the level of genus, and the methanogen data were parsed out of the full data set by taxonomy. It was manually determined which 5 methanogenic genera were most active in the D1 (0-5 cm), D3 (10-15 cm), and D6 (20-30 cm) samples independent of time. The genus-summed mean total transcription of these 5 methanogenic genera over time was plotted in R using ggplot^66^. To represent the activity of individual MAGs over time and depth, the mean MAG-level summed geTMM scores were plotted as a heatmap using ggplot in R.

### Variable Importance (VIP) scores

Variable importance scores (VIP) are used to estimate a variables contribution to PLS regression, with predictors assigned high scores considered important for the PLS prediction of the tested response variable. Here, VIP were calculated as per Chong et al.^81^ in R to correlate methanogen MAG activity – or genome expression- and field methane data. To generate methane data, a numerical model was used to combine chamber and peeper measurements to determine the rates of methane production as outlined in Angle et al^16^. For MAG activity, the aforementioned summed average MAG activity table (see above) was used. Significant VIP scores (>2) were plotted using ggplot in R.

## Supporting information

Supplemental Table 1. Complete list of samples included in the Multi-Omics For Understanding Climate Change (MUCC) v2.0.0 database. Supplemental Tabl

## Acknowledgements

This work was partially supported by awards from the National Science Foundation (NSF) grants EAR-2029686 (EKB, MJW), DEB-1754756 (JEK) and PRFB-2109592 (AMO) and U.S. Department of Energy (DOE) Office of Science, Office of Biological and Environmental Research (BER) grants DE-SC0023084 (EKB, JBE, DKMF, SB, EJW, GB, MAB, KCW, MJW), DE-SC0021350 (MAB, KCW), *DE-SC000054* (DKMF, EJW, GB, KCW), DE-SC0023456 (VXR, RKV) and contributions by JEK were funded by DE-SC0007144, DE SC0012088, and DE-SC0023297. Metagenomic and metatranscriptomic sequencing was performed at the Joint Genome Institute under a Community Science Program and the University of Colorado Anschutz’s Genomics Shared Resource. The work (proposal ID: 504205, 505780) conducted by the U.S. Department of Energy Joint Genome Institute (https://ror.org/04xm1d337), a DOE Office of Science User Facility, is supported by the Office of Science of the U.S. Department of Energy operated under Contract No. DE-AC02-05CH11231. Work conducted at the Genomics Shared Resource at University of Colorado was supported by the Cancer Center Support Grant (P30CA046934).

## Supplementary Information

**Figure S1.**
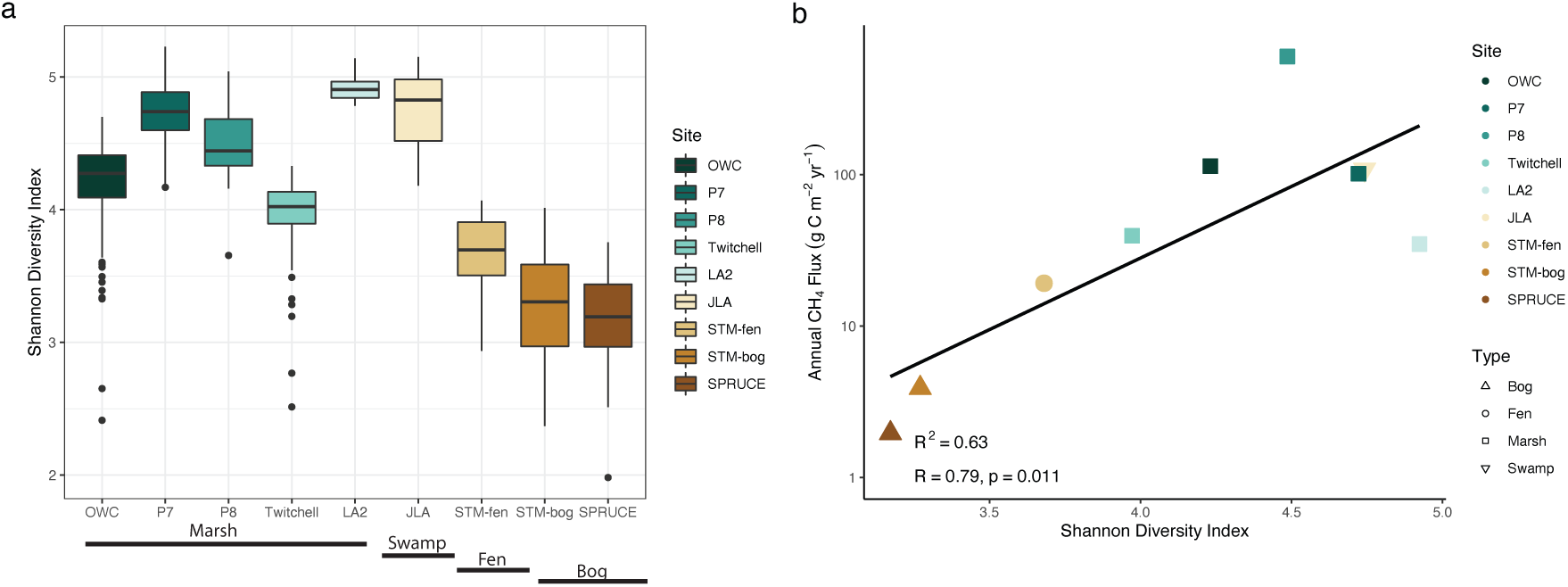
(A) Alpha diversity measured using the Shannon diversity index of the total community was significantly higher in swamps and marshes compared to bog and fen (p<0.001). (B) Total community alpha diversity was significantly correlated with annual CH_4_ flux.

**Figure S2.**
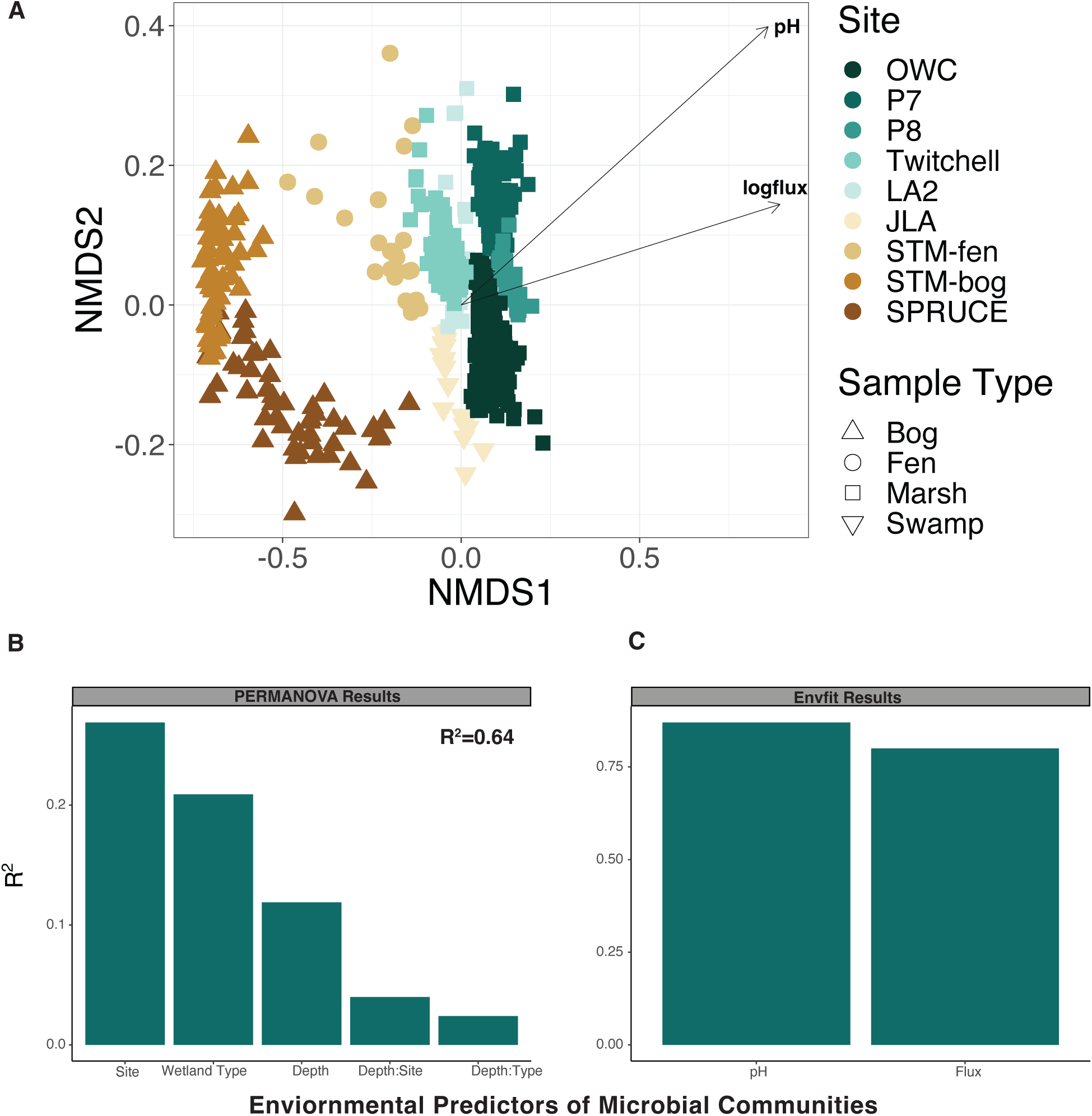
Wetland type and pH are dominant drivers of microbial communities. (A) Nonmetric multidimensional scaling (NMDS) oridination of wetland communities was overlayed with significant environmental variables using envfit. (B) Barplot visualizing the relative importance of environmental factors that explain variation in the microbial communities. (C) Higher pH and CH_4_ flux were correlated with microbial communities from marsh sites.

**Figure S3.**
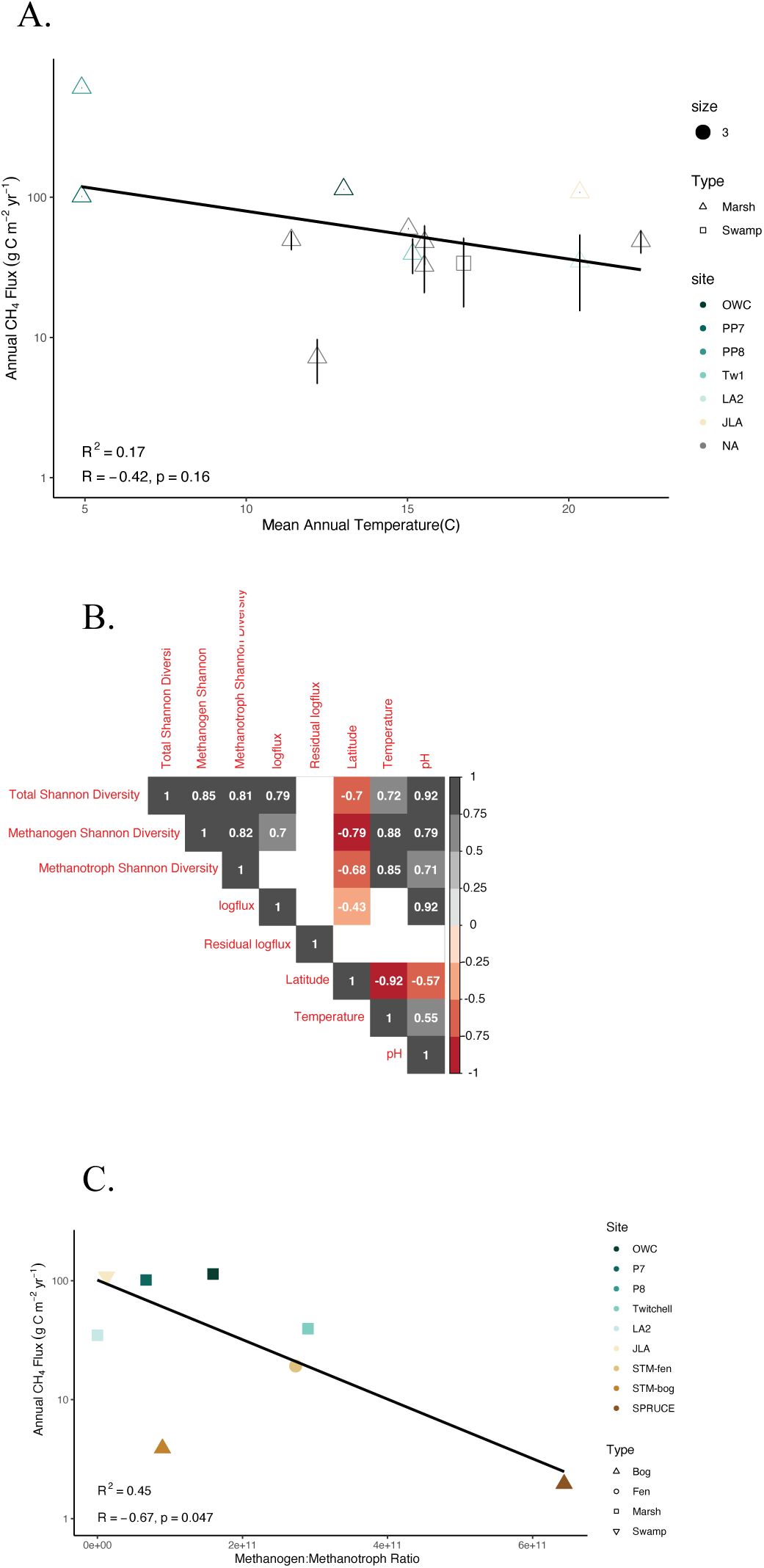
(A) Temperature alone has relatively low predictive power of CH_4_ flux in marshes and swamps. Mean annual temperature of all marsh and swamp found in Delwiche et al. (2021) and in our dataset were compared annual CH_4_ flux. (B) Correlation plot comparing environmental variables to CH_4_ flux and Shannon diversity. White boxes with no value indicates no significant correlation between variables. (C) The ratio of methanogen to methanotroph relative abundance is significantly correlated to CH_4_ flux.

**Figure S4.**
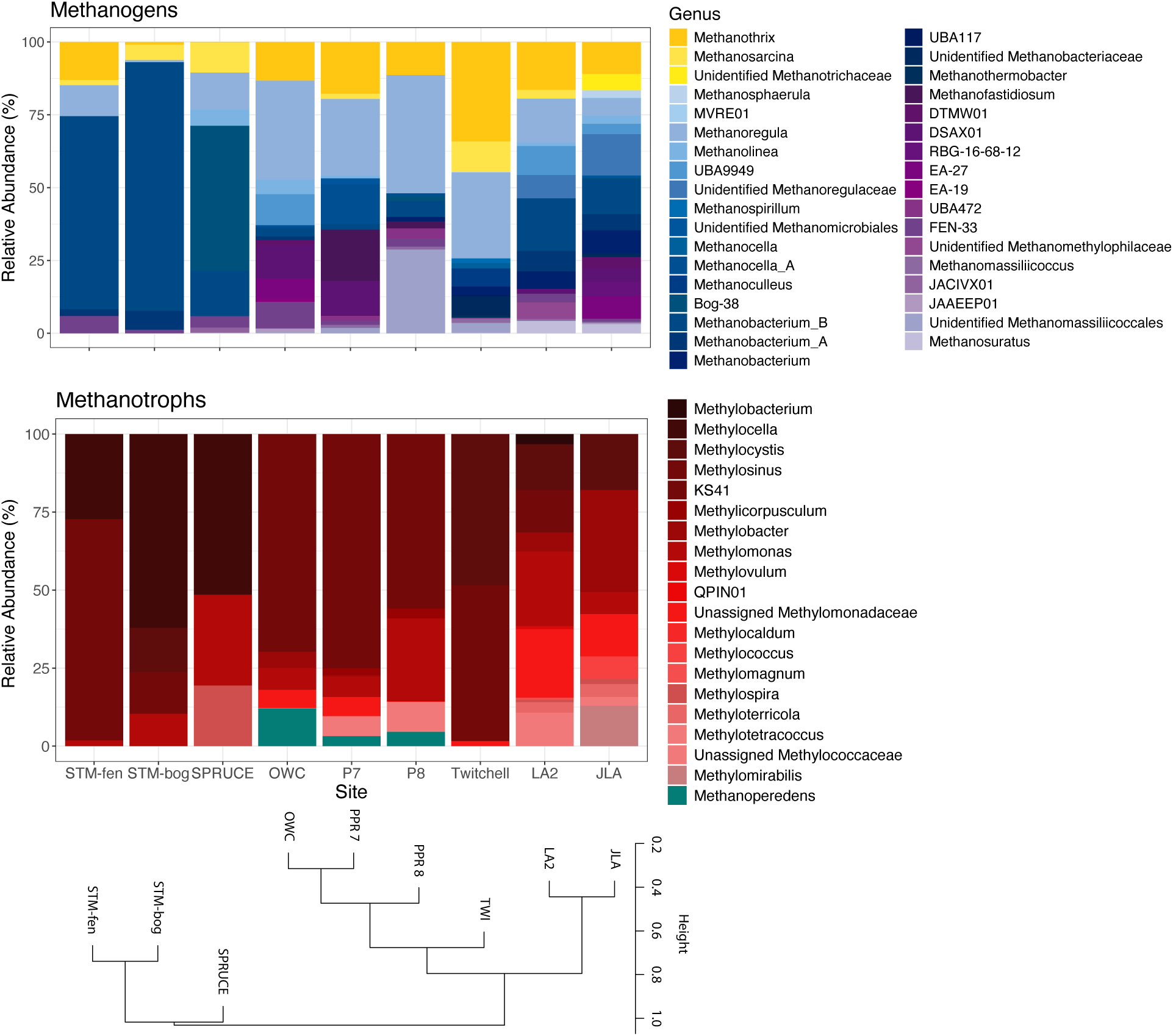
Across 9 wetlands, the highest methane emitting wetlands (PPR P7, OWC) shared similar methane microbial communities despite differences in geography. Amplicon taxonomic data were mined for known methanogen or methanotroph membership and relative abundance. Dominant and prevalent methanogen genera include *Methanoregula* and *Methanothrix*. Methanogens are colored by pathway where acetoclastic methanogens are given in yellow, hydrogenotrophic methanogens in blue and methylotrophic methanogens in purple. Aerobic methanotrophs are given in red while anaerobic methanotrophs are given in teal.

**Figure S5.**
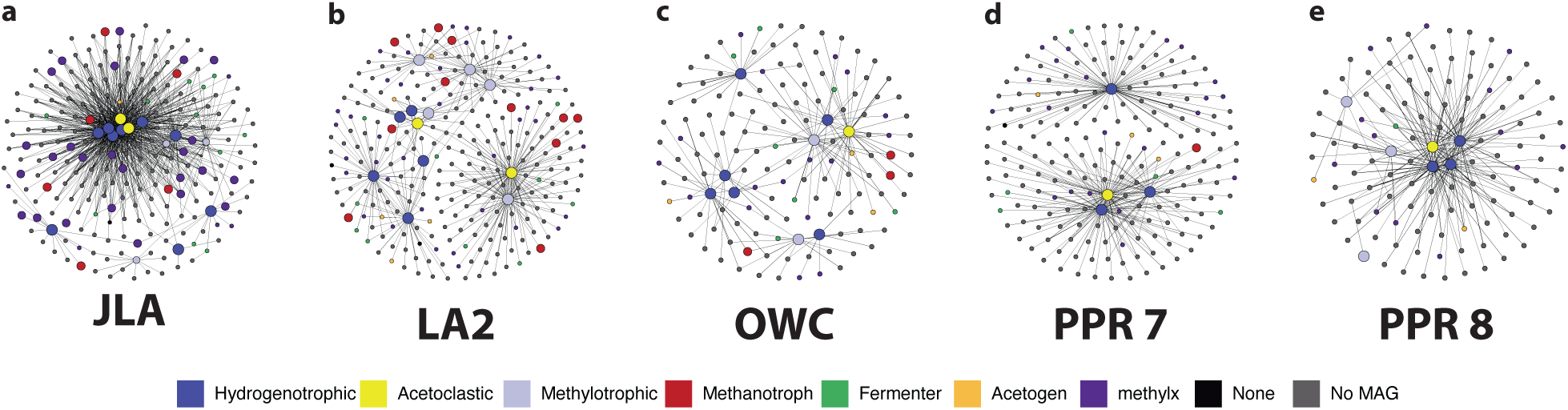
16S rRNA gene co-occurrence networks at each site were built for the methanogen community. Networks were constructed to comprise all significant co-occurrences that included a methanogen.

**Figure S6.**
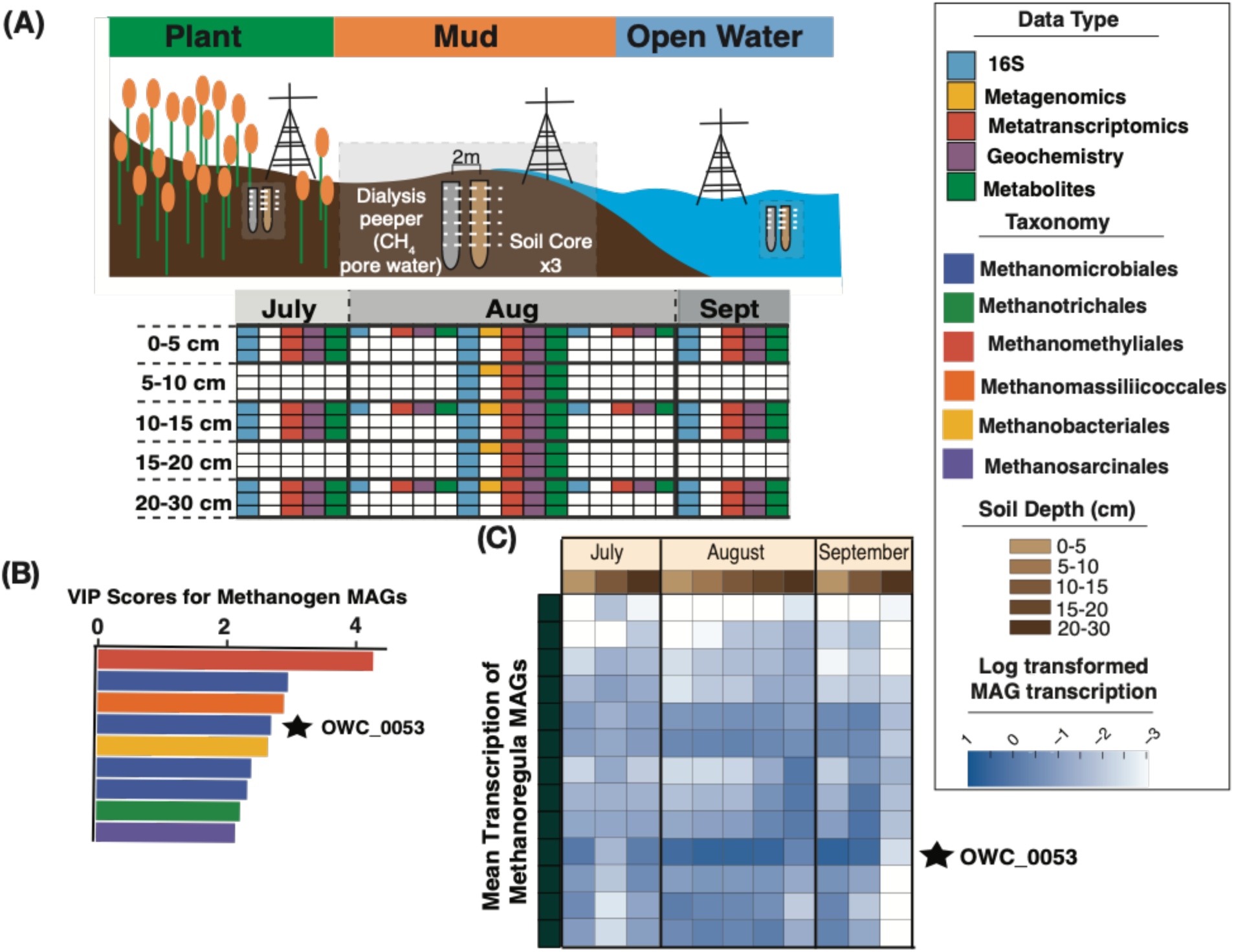
(A) Illustration representing our 2018 sampling campaign in OWC, here used as a case study for exploring the significance of *Methanoregula* in a single wetland. (B) Significant VIP scores (>2) for methanogen MAGs found predictive of CH_4_ fluxes in OWC, including a member of the *Methanoregula* (starred, OWC_0053). (B) Mean transcription of *Methanoregula* MAGs across time and depth in OWC in the mud site type. Members of the genus are active across the site, with the most active MAG representing the sole one found to be significantly predictive of CH_4_ fluxes.

## Notes

https://zenodo.org/records/10822869

### Competing Interest Statement

The authors have declared no competing interest.

